# Phase precession of spindle-slow oscillation coupling across the human brain

**DOI:** 10.1101/2025.11.20.689541

**Authors:** Alex C. Bender, Madeline Taubkin, Mia Y. Bothwell, Kyle Pellerin, Milena K. Pavlova, Rani A. Sarkis, Sydney S. Cash, Alice D. Lam

## Abstract

Spindles and slow oscillations (SO) are fundamental elements of the NREM sleep microarchitecture, often co-occurring in a phase-dependent manner, and this cross-frequency coupling is critical for the temporal coordination of neural activity in sleep. However, spindles and SO occur at different times in different regions, and it is unclear how the coupling of these oscillations is organized across the brain. Here, we provide evidence in humans for a novel spatiotemporal organization of spindle-SO coupling, characterized by a *precession* of the SO phase of spindles along the brain’s anterior-posterior axis. We show that this phase precession relationship is a robust phenomenon and can be quantified across individual subjects. Moreover, the integrity of phase precession strength and slope declines with advancing age. These findings provide new insight into the temporal coordination of sleep rhythms across brain space, linking this coordination to a canonical principle of neural coding.

## INTRODUCTION

Neural oscillations are central to brain function, and understanding how these oscillations are organized across the brain is of fundamental importance in neuroscience. Spanning local to global neural circuits, oscillations support information processing throughout the brain.^1^ The dynamic “up” and “down” states of brain oscillations create temporal windows for the coordinated firing of neuronal assemblies, for spike-timing dependent plasticity, and for inter-regional communication. Oscillations of different frequencies demonstrate hierarchical organization via cross-frequency coupling, exemplified by the phase-amplitude coupling of theta-gamma oscillations in the awake state,^2^ and by phase coupling between slow oscillations (SO) and spindles in sleep.^3^

Spindles and SO are specific oscillatory features of the non-rapid eye movement (NREM) sleep micro-architecture. Spindles (brief 11-16 Hz oscillations) are generated by thalamocortical networks,^4^ and are modulated by neocortical SO (0.5-1.5 Hz), preferentially occurring at specific phases of the SO. This coupling of spindles with SO is integral to the active systems consolidation theory of memory.^5–7^ Sleep spindles and SO are also associated with general cognition,^4,8^ brain health^9^ and aging,^10^ and both are disrupted in brain disorders, such as epilepsy,^11,12^ schizophrenia,^13^ and Alzheimer’s disease.^14^ However, because spindles and SO occur locally at different times and in different regions^15–17^, a fundamental unanswered question is whether and how the coupling of these rhythms is organized across the brain.

To better understand the whole brain organization of spindle-SO coupling, we leveraged a human dataset of full scalp EEGs recorded overnight in either the inpatient Epilepsy Monitoring Unit (EMU) or the home/ambulatory setting (AMB). We then examined the specific phase relationship between spindles and SO across the brain surface from the most anterior to posterior positions, hypothesizing that the SO phase preference of spindles would be non-uniform across brain space, and instead demonstrate a systematic relationship with position along the cortical surface.

## RESULTS

### Phase-position relationship of spindle-SO coupling

The dataset consisted of 101 participants (eTable 1) free of major neurological disease, ranging in age from 20 to 85 years (mean 57 years +/- std 19.2, 34% male), with scalp EEGs recorded overnight in either the inpatient EMU ( N=58) or the home (AMB) setting (N=43). Initial observations were made in the EMU group, which consisted of patients who underwent evaluation for spells in the EMU and were determined to not have epilepsy. We examined how spindle-SO phase coupling varied across the brain surface, as follows (Fig. 1a-d): Overnight scalp EEG was pre-processed and automatically sleep staged.^18^ For each participant and at each EEG channel, spindles were detected during N2 and N3 sleep stages (eFig. 1), and the SO phases corresponding to the spindle peaks were extracted to compute the mean SO phase preference of spindles. The mean phases for each participant were then included in a group-level circular plot and a mean phase was determined at each channel. Along the anterior-posterior axis, we observed a clear pattern of spindle-SO phase preferences, with a systematic shift to earlier phases at each subsequent position. In other words, spindles exhibited SO *phase precession* relative to their recorded position along the brain’s anterior-posterior axis (Fig. 1e-f).

**Figure 1.**
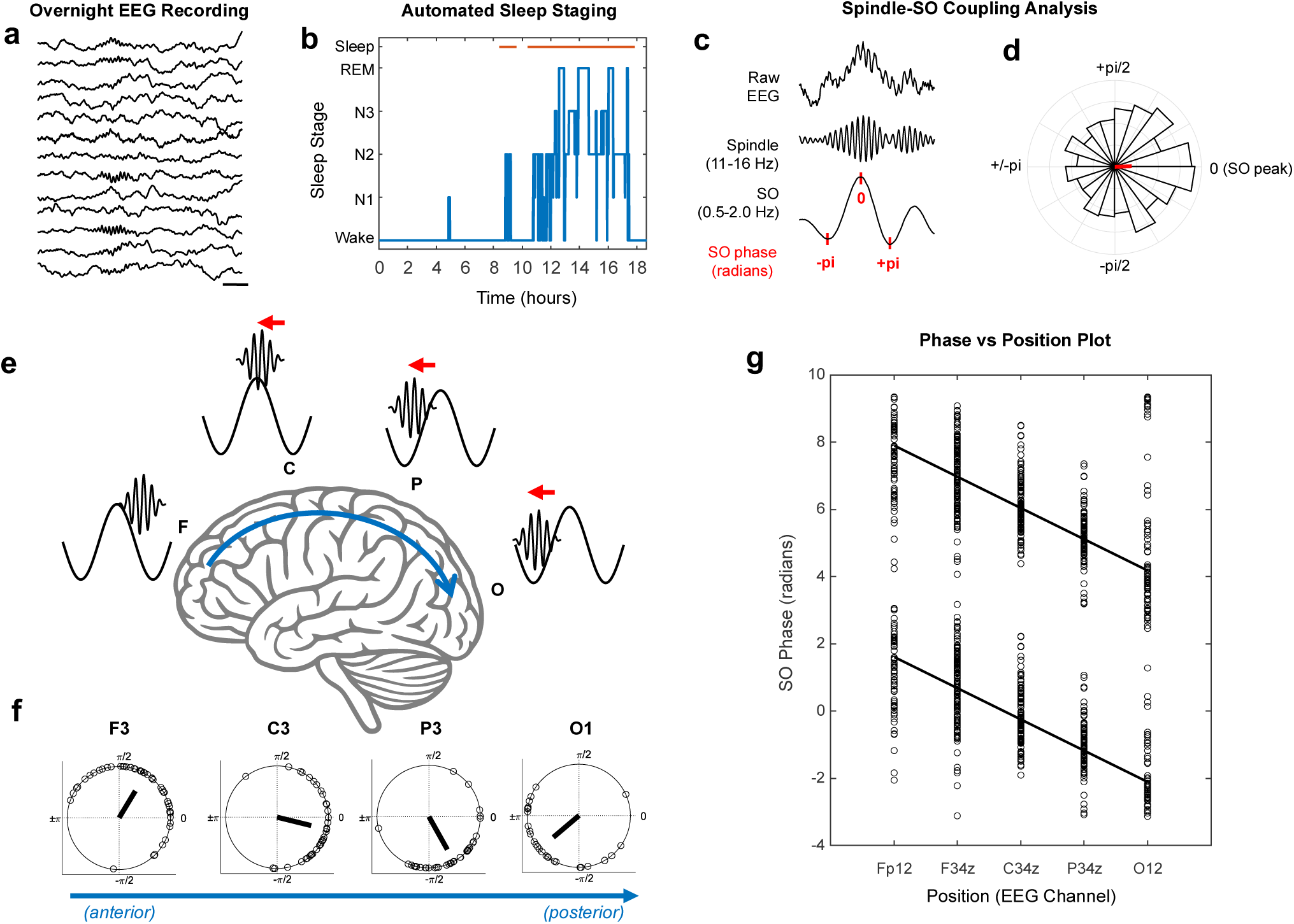
Phase-position relationship of spindle-SO coupling across the brain. **a**, Example EEG tracing recorded from a single subject in NREM sleep. Scale bar equals 500 ms. **b**, Example of sleep hypnogram from the same subject derived from automated sleep-staging of the EEG. **c**, Example of spindle-SO coupling from the same subject, with raw EEG tracing (top), filtered spindle activity (middle), and corresponding filtered SO signal (bottom) with the SO phases indicated in red. **d**, Circular phase histogram representing the SO phases of spindles for all N2 and N3 sleep for the subject in panels a-c. The red bar indicates the mean phase preference, occurring near the SO peak, or at a phase of ∼0 radians. **e-f**, Advancement of the SO phase of spindles from anterior to posterior positions over the brain’s surface. In panel **f**, the mean SO phase is shown for all EMU participants (N=58, each circle = 1 subject) in a circular plot for the labeled EEG channels, F3, C3, P3, and O1. Panel **e** shows an illustration of the SO phases of spindles corresponding to the mean phases in **f**. **g**, A phase-position plot showing the mean SO phase for all EMU participants (N=58) relative to EEG channel position. The data are plotted twice for each phase (phase+0 and phase+2pi), for better visualization on a linear axis. A “best fit” regression line is shown (p<0.0001, R=0.62, Slope=-0.93).

This *phase precession* relationship of spindle-SO coupling was visualized in a phase-position plot (Fig 1g). We then used circular statistics to quantify the slope^19^ and strength (R-value) of the phase precession relationship. In EMU participants, the mean slope was approximately -0.9 radians, or -51.5 degrees, of phase shift for every 1 unit change in electrode position (p<0.0001, R=0.62), which corresponds to 20% of the distance measured from the nasion to inion in a standard 10-20 EEG system.

### Sensitivity analyses of phase precession

To examine whether the observed phase precession relationship was sensitive to the choice of the EEG reference electrode, montage, or recording environment, we re-referenced the EEGs from EMU participants and compared the phase-position plots derived from a common average reference montage, a neck (Cs2) referential montage, and a longitudinal bipolar montage (Fig. 2a-c). Phase precession was evident regardless of the reference channel used (p<0.0001 in all plots). The common average reference and longitudinal bipolar montages produced phase-position relationships with similar slopes (-0.93 vs - 0.90) and R-values (0.62 and 0.57), whereas the Cs2 referential montage yielded a smaller, but still highly significant, slope (-0.51, p<0.0001), likely related to a reduced phase shift at the occipital channels, which were in close proximity to the Cs2 reference.

**Figure 2.**
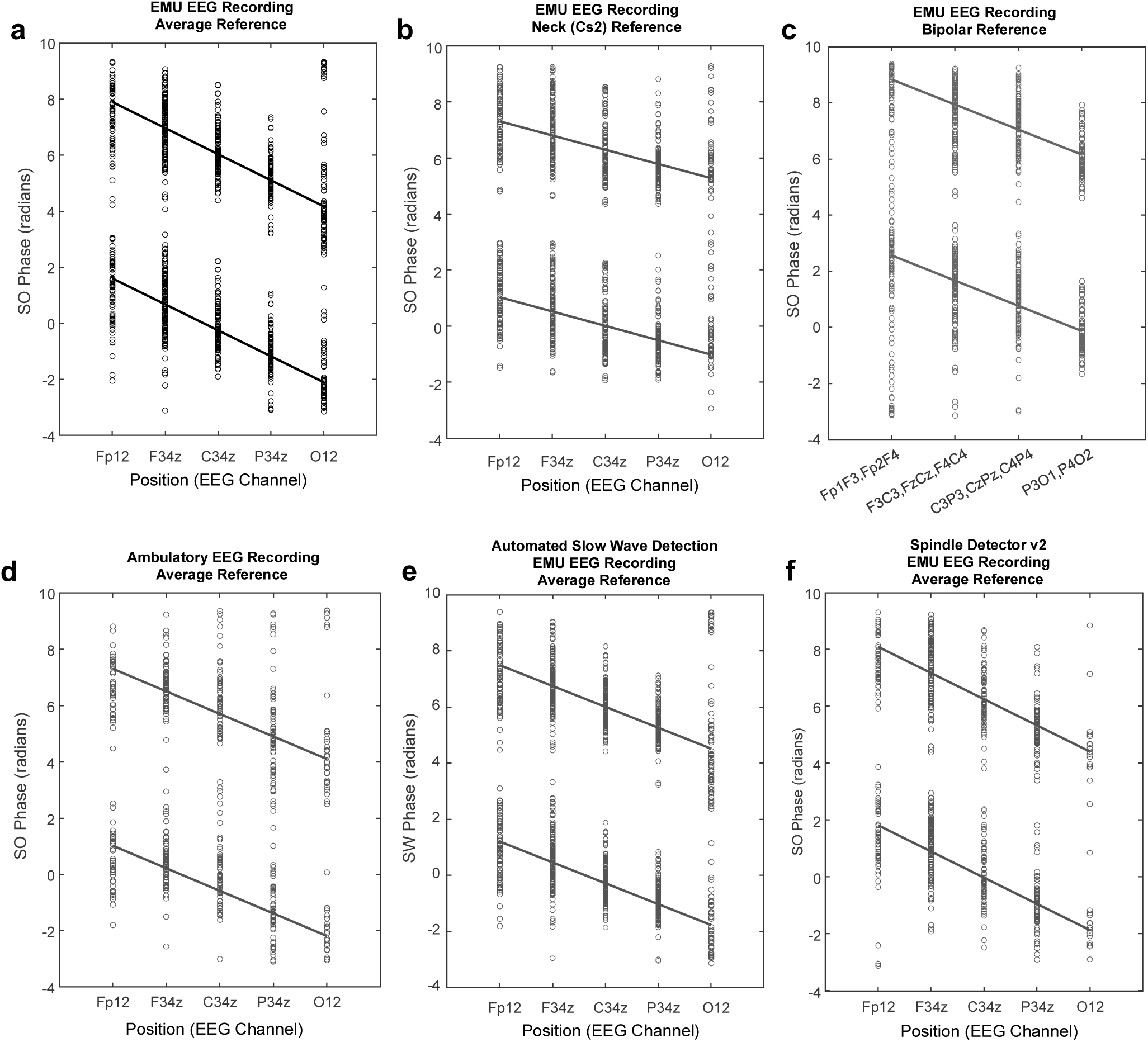
Phase precession of spindle-SO coupling is a robust pheno enon. Phase-position plots are shown for EMU participants (N=58; panels **a-c,e,f**) and AMB participants (N=43; panel **d**), as described in Fig. 1g, but with varying methodology in its measurement. Comparing **a**-**c**, data is shown for EMU participants, varying the EEG reference and montage. **a**, Common average reference (p<0.0001, R=0.62, Slope=-0.93). **b**, Neck (Cs2) reference (p<0.0001, R=0.50, Slope=-0.51). **c**, Longitudinal bipolar reference (p<0.0001, R=0.57, Slope=-0.90). Comparing **a** and **d**, the common average reference is used in both, but two different cohorts recorded in different environments are shown: **a**, EMU participants and **d**, AMB participants (p<0.0001, R=0.65, Slope=-0.80). Comparing **a** and **e**, the common average reference is used in both, but the SO phase is derived from either: **a**, the continuous filtered SO signal, or **e**, discrete slow wave detections (p<0.0001, R=0.60, Slope=-0.75). Comparing **a** and **f**, the common average reference is used in both, but in **f**, a different spindle detection algorithm is used (p<0.0001, R=0.65, Slope=-0.92).

To examine whether the phase precession relationship was sensitive to the EEG recording environment or equipment, we analyzed an independent dataset of ambulatory EEGs (AMB) recorded in the home setting and in a different cohort of participants. We again observed phase precession (Fig. 2d) with similar characteristics (slope=-0.80, R=0.64, p<0.0001) compared to the EMU dataset.

We then tested whether phase precession depends on the methodology used to derive SO phase values (i.e., whether phase was derived from SO as a continuous signal vs. discrete slow wave events). We automatically detected slow wave events (SW) using previously published algorithms,^10,15^ and extracted the SO phase of spindles during SW detections. We again observed a significant phase precession relationship (Fig. 2e, p<0.0001) with similar characteristics (slope=-0.74, R=0.60) compared to when phases were extracted from the continuous SO signal (slope=-0.93, R=0.62). Finally, we tested whether phase precession was sensitive to the methodology of spindle detection by comparing our spindle detector to a previously published algorithm.^13^ The phase precession relationship remained highly significant (Fig. 2f, p<0.0001) and with similar characteristics (slope=-0.92, R=0.65). In summary, we found that phase precession was a robust phenomenon and was observed with similar characteristics across different montages, recording environments, and spindle and SO detection algorithms.

### Phase precession analysis across individuals

To examine how phase precession differs across individuals, we constructed phase-position plots for each subject, using SO phase values for all spindles over each subject’s EEG recording. We computed the best fit line and the phase precession slope from these raw phase values (Fig. 3). The majority (84.1%) of all participants exhibited a significant phase precession relationship, with a mean slope of - 0.87 (+/- 0.24) radians per 1 unit change in electrode position. Examples of individuals with strong phase precession are shown in Fig. 3a-c. However, a small proportion of individuals showed weak or no phase precession (Fig. 3d-f).

**Figure 3.**
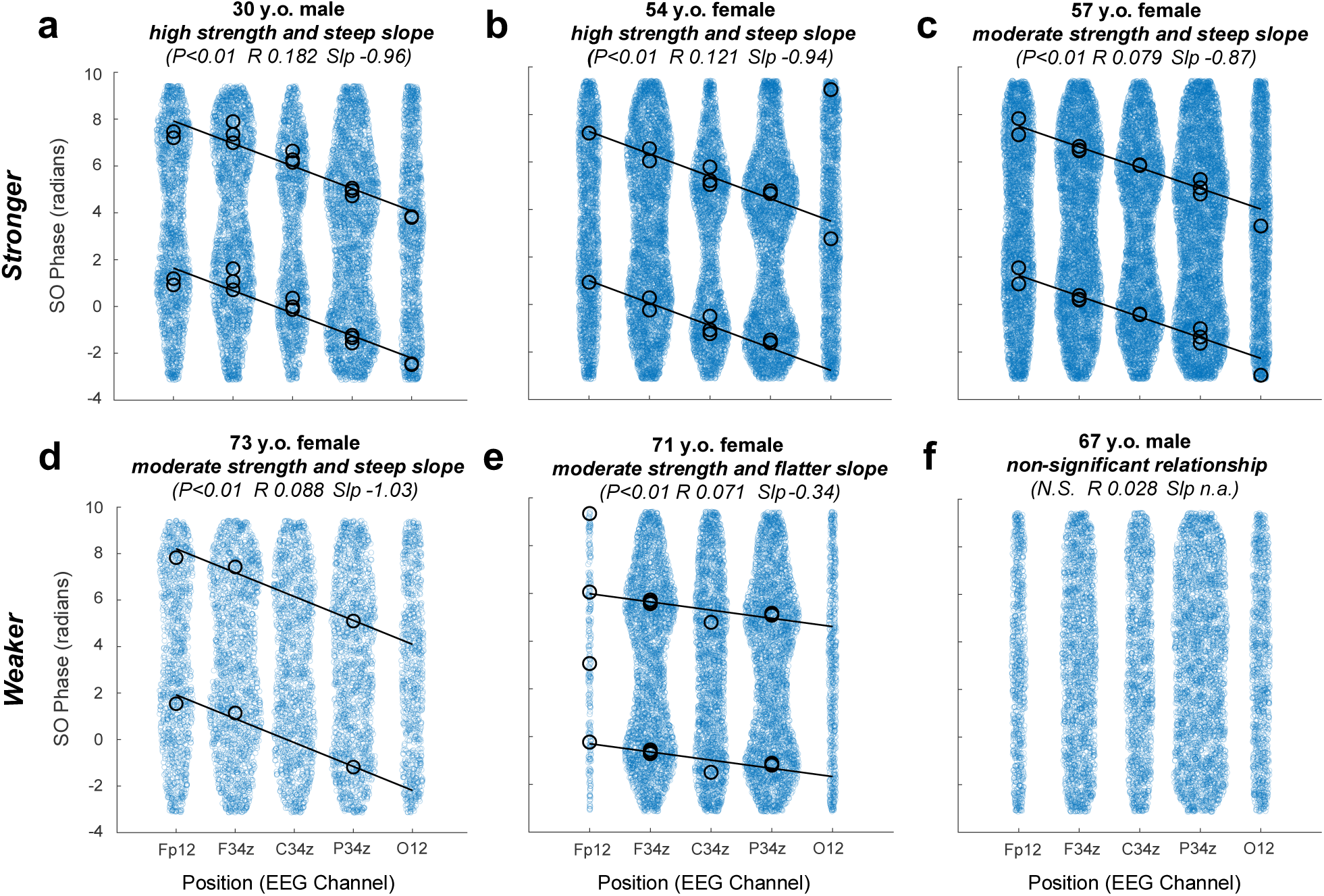
Analysis of phase precession across individual subjects. Phase-position plots are shown for 6 different exemplar subjects, **a-f**, with phase precession relationships ranging from high strength (higher R value) and steep slope (more negative) to non-significant. Blue circles represent each SO phase value for all spindles recorded over N2 and N3 sleep. The regression line is computed from these data. Black circles represent the mean phase at each EEG channel and are only shown when there was a significant non-uniform distribution of the phases. The subject demographics (age and sex) and circular-linear statistics for the phase-position relationship are shown above each plot (Slp=slope, N.S.=not significant).

Given the observed inter-individual variability in phase precession properties, we next asked whether this variation could be related to demographic variables, age and sex. Data from all participants were combined for this analysis (sub-group analyses are shown in eFig. 2). We found that older age is associated with reduced phase precession strength (Fig. 4a; F=5.15, b=-0.001, R=0.37, p=0.008, adjusting for total spindle count and group). Consistent with this, there was a significantly lower proportion of participants in the oldest quartile who exhibited significant phase precession compared to those in the youngest quartile (71.9% vs 96.0%, chi^2^=5.65, p=0.018). We also observed a significant association of age with the phase precession slope, with older ages having a flatter slope (Fig. 4c; F=2.64, b=-0.005, R=0.23, p=0.016, adjusting for total spindle count and group). In contrast, we did not find any significant differences between male and female sex for phase precession strength (Fig. 4b; F=2.49, p=0.64) or slope (Fig. 4d; F=0.77, p=0.53). Thus, advancing age is associated with a reduction in the strength and a flattening of the slope of phase precession.

**Figure 4.**
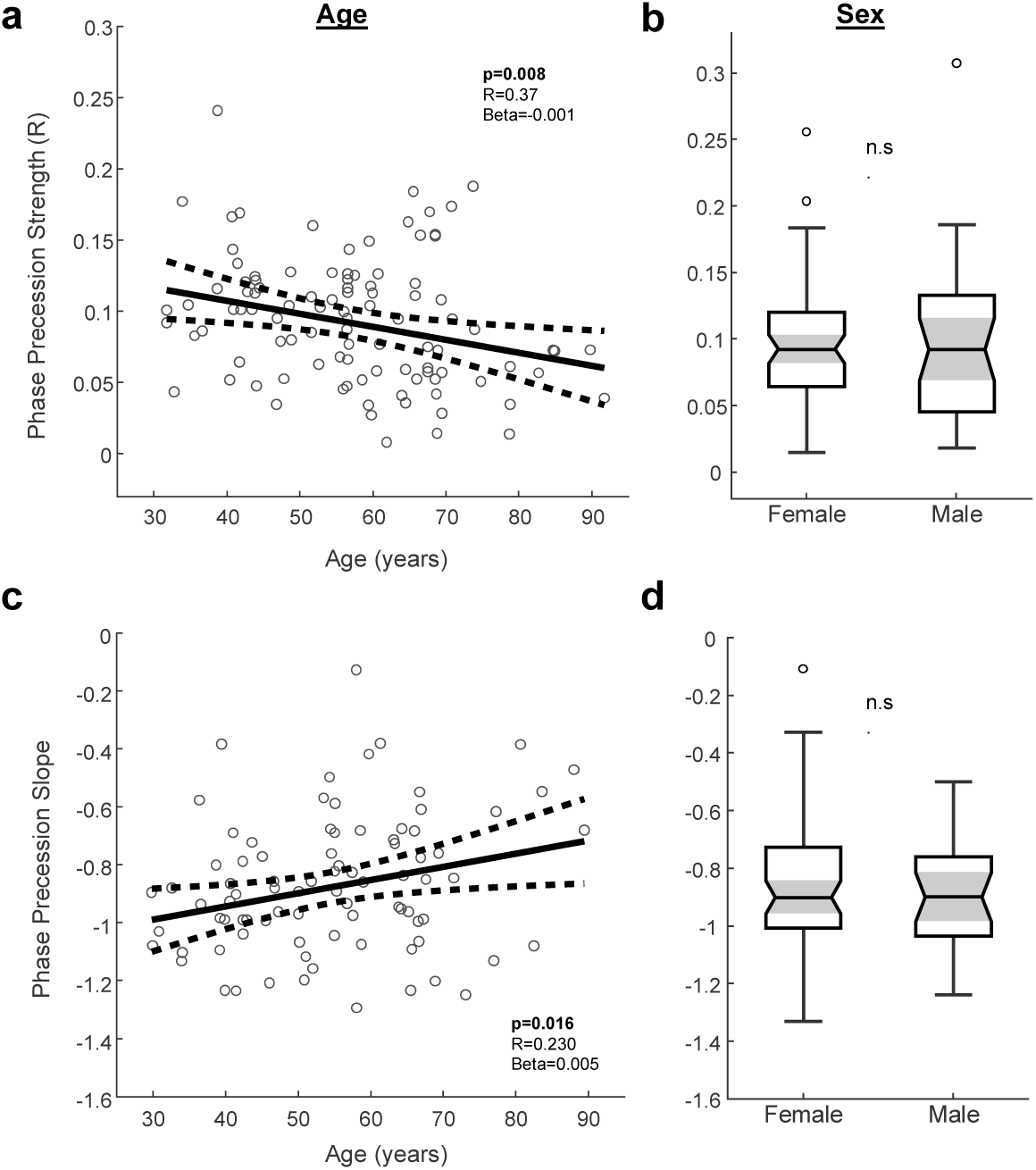
Phase precession properties are associated ith age but not sex. Data are shown for all participants (EMU + AMB, N=101). **a,** Scatter plot of model-adjusted phase precession strength versus age. There was a significant negative association, indicating reduced strength of phase precession with advancing age (F=5.15, b=-0.001, R=0.37, p=0.008, adjusting for total spindle count and group). **b**, Phase precession strength was not significantly different between female and male participants (data shown in boxplots, p>0.05). **c**, Scatter plot of model-adjusted phase precession slope versus age. There was a significant positive association, indicating a flattening of the slope with advancing age (F=2.64, b=-0.005, R=0.23, p=0.016, adjusting for total spindle count and group). **d**, Phase precession slope was not significantly different between female and male participants (p>0.05).

## DISCUSSION

In summary, we report a novel phase-position relationship of spindle-SO coupling observed with human scalp EEG, characterized by a systematic *precession* of the SO phase of spindles along the anterior-posterior axis of the brain. We show that this is a robust phenomenon and can be quantified across individual subjects. Moreover, the integrity of phase precession strength and slope declines with advancing age.

The concept of *phase precession* is fundamental to neural coding, having first been used by O’Keefe and Recce in 1993 to describe the firing of hippocampal place cells with respect to the local theta oscillations.^20^ In the three decades since, theta phase precession has been demonstrated for other cell types, such as grid cells,^21^ and in other animal species, including humans.^22^ However, it has never been reported for sleep oscillations or across brain regions. We propose that phase precession of spindle-SO coupling represents a fundamental spatiotemporal organization of these oscillatory elements across the brain, and that its specific phase relationship may be an important coding mechanism for how information is processed in sleep.

Spindles and SO have received substantial attention for their roles in memory consolidation. The coupling of these oscillations, along with ripples, regulates the communication between the hippocampus and neocortex,^6,7^ facilitating the formation of long-term memory representations.^5–7^ However, spindles and SO occur locally at different times in different regions. Moreover, different brain regions host different aspects of a given experience that must be bound together into a unique memory representation.^15–17^ How this is all coordinated across the brain in sleep is unknown. Phase precession may offer a potential solution by providing a phase code for brain space that could facilitate the binding of multi-modal information as memory traces are reactivated in this complex hippocampal-neocortical dialogue.

Finally, our observations raise several new and important questions that warrant further investigation, particularly regarding the neural mechanisms responsible for generating phase precession of sleep oscillations, how this phenomenon fits into current theories on the neuronal and computational underpinnings of memory and cognition, and whether and how phase precession is altered in neurological disease.

## METHODS

### Study population

This was a retrospective analysis of data collected from a total of 101 participants with overnight full scalp EEG recorded between 2008 and 2024. EEGs were recorded either in the home setting (AMB) or in the Epilepsy Monitoring Unit (EMU) at Massachusetts General Hospital (MGH) or Brigham and Women’s Hospital (BWH). The dataset included 101 participants without major neurological disease, ranging in age from 20 to 85 years. 43 participants had ambulatory EEG recorded in the home setting (AMB), collected as part of an aging and Alzheimer’s disease study and were determined to be cognitively unimpaired.^23^ The other 58 participants had EEG recorded in the EMU for evaluation of spells and were determined to not have epilepsy based on the following criteria: 1) the absence of ictal EEG correlate, 2) absence of inter-ictal epileptiform abnormalities, and 3) clinical judgement of the EMU team or outpatient epileptologist that the patient did not have epilepsy. Patients were excluded if there was a history of prior brain surgery, large territory stroke or high-grade tumor. Demographic and clinical data were obtained from electronic medical records.

### EEG Recording and Sleep Staging

All participants underwent overnight EEG recordings with placement of full scalp EEG electrodes according to the International 10-20 system. Ambulatory scalp EEG recordings were obtained with participants sleeping in their home environment, while EMU recordings were obtained with participants sleeping in a hospital bed. Continuous EEG recordings were acquired at a minimum of 200 Hz with Xltek hardware (Natus Medical Inc., Middleton, WI). EEG data were down-sampled to 128 Hz, re-referenced using a common average reference, or as otherwise noted in the main text, and bandpass filtered between 0.1 to 50 Hz. EEG recordings were pre-processed offline using Matlab 2022a (Mathworks, Inc. Natick, MA), the FieldTrip toolbox^24^, and custom scripts.

Sleep staging was performed in Python using an automated deep learning algorithm that was previously validated in our lab (https://github.com/mauriceaj/CRNNeeg-sleep).^18^ The algorithm scores the EEG in non-overlapping 30-second epochs and labels each epoch as N1, N2, N3, REM or awake. For participants with more than one night of EEG, only the first 24 hours was used.

### Analysis of Spindles and SO

Pre-processed EEGs were analyzed using custom scripts and toolboxes (signal processing, statistics and wavelet) in Matlab. Prior to analysis, all EEGs were visually inspected to confirm consistent polarity by ensuring that eye blink deflections in awake segments were upward. Analysis was subsequently restricted to artifact-free EEG during stages N2 and N3 sleep. Automated artifact rejection was applied to 5-second windows based on 3 criteria: 1) excessively high voltages (> 400 uV), 2) excessively low voltage ranges (standard deviation below 0.01 uV), or 3) excessive myogenic artifact (standard deviation of the epoch’s 30-50 Hz bandpass filtered EEG signal exceeded 3 times the standard deviation of the whole EEG’s 30-50 Hz bandpass filtered EEG signal). Five-second windows that met any of the 3 criteria above were removed from all analyses.

Spindles were detected automatically (https://github.com/acbender/sleep), based on amplitude and duration thresholds as described previously.^11^ Briefly, the wavelet-derived power of the EEG signal in the spindle frequency range (11-16 Hz) was computed and smoothed with a 100 ms moving average window. The smoothed signal was converted to z-scores, and spindle events were identified when the z-scored signal exceeded 2 for at least 300 ms and exceeded 3 at its maximum within the putative spindle event. An alternative spindle detector was used in the analysis for Fig. 2f, following the methods previously published by Wamsley et al., 2012.^13^

To analyze coupling of spindles to the slow oscillation (SO), the Hilbert transform was used to compute the instantaneous phase angles of the filtered SO signal (0.5-2 Hz). SO phases corresponding to the times of spindle events, taken at the spindle peak, were extracted and saved for each EEG channel that was used in the phase precession analysis. Circular statistics were used to compute the mean phase angle (i.e. phase preference) over the duration of the overnight EEG recording for each channel, and analysis of the mean phase was restricted to when there was a significant non-uniform distribution to the phases by the Rayleigh test.

In sensitivity analyses, spindle-SO coupling was additionally examined by extracting discrete slow wave (SW) events, as opposed to considering the SO as a continuous signal. The SW events were detected by an automated algorithm that is based on previously published reports.^10,15^ Briefly, after applying a zero-phase, 0.5 to 4.0 Hz bandpass filter to the artifact-free EEG during N2 and N3 sleep, negative deflections were detected and only waves with consecutive zero crossings separated by 0.25 to 1.0 seconds (0.5 to 2.0 Hz) and with a peak in envelope amplitude above the 75^th^ percentile of the filtered signal were considered. The SW phase corresponding to the peaks of co-occurring spindle events was then extracted as described above.

### Visualization and Quantification of Phase Precession

Phase-position plots were constructed by considering each of the parasagittal EEG electrodes (Fp1, Fp2, F3, Fz, F4, C3, Cz, C4, P3, Pz, P4, O1, O2) as representing a relative position along the scalp in the anterior-to-posterior direction. In the standard 10-20 system, the EEG technician places each of the frontopolar, frontal, central, parietal, and occipital electrodes equidistant from each other along the anterior-posterior axis (each inter-electrode spacing represents 20% of the distance from the nasion to inion). Thus, each grouping of electrodes (e.g. Fp1 and Fp2) was considered together and was represented as 1 unit in distance from the next grouping (x-axis). The phase values (y-axis) were the SO or SW phases corresponding to the spindle peaks, as detailed above. To better visualize circular values falling at the extremes on a linear scale, the phases were plotted twice, with the duplicative values equal to the original values + 2*pi (a common convention^19,22^).

To quantify the slope and best fit line of the phase precession relationship, we adopted the methods established by Kempter, et al.^19^ In contrast to other methods that treat the circular variable as linear and restrict the phases to a predefined range, this approach does not set arbitrary cutoffs or require *a priori* knowledge of the data. Instead, a range of reasonable slope values is tested and the best fit line is the result that minimizes the sum of circular distances between the data and the line (equivalent to maximizing the mean resultant length of the residuals).^19^ To quantify the correlation coefficient and the statistical significance of this circular-linear relationship, we used a circular-linear correlation function developed by Berens et al.,^25^ available in the Circular Statistics Toolbox in Matlab. The phase precession relationship was considered significant when the p-value was less than 0.05 and the slope was non-zero.

To plot and quantify phase precession at the group level, we used the mean phase values for each subject at each electrode (as in Fig. 1g). This was calculated from all spindle-SO coupling events over the duration of the subject’s EEG recording for each electrode, and analysis was restricted to when there was a significant non-uniform distribution to the phases by the Rayleigh test at p<0.05 (i.e. there was a significant phase preference). In these plots, each electrode for each subject is represented as a single mean phase value. To plot and quantify phase precession at the individual subject level (as in Fig. 3), we used the raw phase values for all spindle-SO coupling events over the duration of the subject’s EEG recording for each electrode. This approach preserves the raw phase data in the phase precession calculation, even when there is not a significant phase preference at a given electrode. As seen in Fig. 2, the regression line still passes through the mean phase values even though the mean phases are not actually used to derive the line. In contrast to the former method, the correlation coefficients (R value) are lower due to the greater spread of the raw phases in the calculation, but the phase precession relationship remains highly significant.

### Sensitivity Analyses

Several sensitivity analyses were done to examine the robustness of the phase precession relationship: 1) To examine phase precession across different recording environments, we compared those subjects with EMU recordings performed in the hospital against those with ambulatory EEG recordings performed at home (AMB). 2) To examine the robustness of the phase precession relationship to the recording montage, we re-referenced the EEG recordings using 3 different montages and then plotted the phase precession relationship for each recording montage using a common average reference, a neck (Cs2) reference, or a longitudinal bipolar reference. In the Cs2 reference, each cortical electrode is referenced to the same Cs2 electrode. In the bipolar reference, each electrode is referenced to a nearest neighboring electrode following the standard conventions for a longitudinal bipolar montage. 3) To examine if phase precession is influenced by whether the SO signal is considered as a continuous variable or extracted only from discrete slow wave (SW) events, we restricted the analysis to phase values taken only from SW detections. The SW detector is described in more detail above. 4) To examine if phase precession is sensitive to the precise algorithm for spindle detection, we repeated the analysis using a different but previously published algorithm^13^, which is more stringent in its amplitude threshold criteria and does not use a Z-scored signal in the computation.

### Statistical Analyses

Statistics were performed in Matlab using the Statistics and Machine Learning Toolbox. Population statistics are reported as the mean ± standard deviation (s.d.), unless specified otherwise. A value of p < 0.05 was considered significant.

To test the association of phase precession properties with demographic variables (age or sex), we performed linear regressions with age or sex as the independent variable and the phase precession slope or R value (circular-linear correlation coefficient) as the dependent variable. To adjust for each recording and subject having a variable rate and count of spindle-SO coupling events, we included the total count of phase values that was used for the phase precession calculation for each subject as a covariate. This was done to account for the possibility that subjects with fewer spindles or spindle-SO coupling events might have weaker phase precession and lower R values for the circular-linear correlation. Both EMU and AMB datasets were combined for this analysis (N=101). When slope was the outcome variable, only those with a statistically significant phase precession relationship were included (N=85; Fig. 4c,d).

### Standard Protocol Approvals and Patient Consent

All study procedures were performed under protocols approved by the Mass General Brigham Institutional Review Boards. Written informed consent was obtained from all prospectively recruited research participants.

### Data and Code Availability

Data that support the findings of this study are available from the corresponding author upon reasonable request, subject to internal review to protect patient confidentiality, and after completion of a data sharing agreement in accordance with Mass General Brigham institutional guidelines. The code for the phase precession analysis is available on GitHub (https://github.com/acbender/sleep).

## ACKNOWLEDGEMENTS

We thank the research participants for contributing to this study. This study was supported by grants from the NIH NINDS (ACB and SSC: UE5NS065743, ACB: K23NS140388; ADL: K23NS101037; RAS: K23NS119798), Mass General Neuroscience Transformative Scholars Award (ACB), Alzheimer’s Association (ADL) and the American Academy of Neurology Institutes (ADL). The Mass ADRC is supported by the NIH (P30AG062421).

## Notes

### Competing Interest Statement

The authors have declared no competing interest.

## REFERENCES

1. Buzsáki, G. & Watson, B. Brain rhythms and neural syntax: implications for efficient coding of cognitive content and neuropsychiatric disease. Dialogues Clin Neurosci 14, 345–367 (2012).

2. Lisman, J. E. & Jensen, O. The Theta-Gamma Neural Code. Neuron 77, 1002–1016 (2013).

3. Staresina, B. P. et al. Hierarchical nesting of slow oscillations, spindles and ripples in the human hippocampus during sleep. Nat Neurosci 18, 1679–1686 (2015).

4. Fernandez, L. M. J. & Lüthi, A. Sleep spindles: Mechanisms and functions. Physiol Rev 100, 805–868 (2020).

5. Rasch, B. & Born, J. About sleep’s role in memory. Physiol Rev 93, 681–766 (2013).

6. Klinzing, J. G., Niethard, N. & Born, J. Mechanisms of systems memory consolidation during sleep. Nat Neurosci 22, 1598–1610 (2019).

7. Staresina, B. P., Niediek, J., Borger, V., Surges, R. & Mormann, F. How coupled slow oscillations, spindles and ripples coordinate neuronal processing and communication during human sleep. Nat Neurosci 26, 1429–1437 (2023).

8. Fogel, S. M. & Smith, C. T. The function of the sleep spindle: A physiological index of intelligence and a mechanism for sleep-dependent memory consolidation. Neuroscience and Biobehavioral Reviews vol. 35 1154–1165 (2011).

9. Sun, H. et al. Brain age from the electroencephalogram of sleep. Neurobiol Aging 74, 112–120 (2019).

10. Helfrich, R. F., Mander, B. A., Jagust, W. J., Knight, R. T. & Walker, M. P. Old Brains Come Uncoupled in Sleep: Slow Wave-Spindle Synchrony, Brain Atrophy, and Forgetting. Neuron 97, 221–230.e4 (2018).

11. Bender, A. C. et al. Altered Sleep Microarchitecture and Cognitive Impairment in Patients with Temporal Lobe Epilepsy. Neurology 101, E2376–E2387 (2023).

12. Sarkis, R. A. et al. Epilepsy and sleep characteristics are associated with diminished 24-h memory retention in older adults with epilepsy. Epilepsia 64, 2771–2780 (2023).

13. Wamsley, E. J. et al. Reduced sleep spindles and spindle coherence in schizophrenia: Mechanisms of impaired memory consolidation? Biol Psychiatry 71, 154–161 (2012).

14. Páez, A., et al. Sleep spindles and slow oscillations predict cognition and biomarkers of neurodegeneration in mild to moderate Alzheimer’s disease. Alzheimer’s and Dementia 10.1002/alz.14424 (2025).

15. Nir, Y. et al. Regional Slow Waves and Spindles in Human Sleep. Neuron 70, 153–169 (2011).

16. Andrillon, T. et al. Sleep spindles in humans: Insights from intracranial EEG and unit recordings. Journal of Neuroscience 31, 17821–17834 (2011).

17. Piantoni, G., Halgren, E. & Cash, S. S. Spatiotemporal characteristics of sleep spindles depend on cortical location. Neuroimage 146, 236–245 (2017).

18. Abou Jaoude, M., et al. Expert-level automated sleep staging of long-term scalp electroencephalography recordings using deep learning. Sleep 43, (2020).

19. Kempter, R., Leibold, C., Buzsáki, G., Diba, K. & Schmidt, R. Quantifying circular-linear associations: Hippocampal phase precession. J Neurosci Methods 207, 113–124 (2012).

20. O’Keefe, J. & Recce, M. L. Phase relationship between hippocampal place units and the EEG theta rhythm. Hippocampus 3, 317–330 (1993).

21. Hafting, T., Fyhn, M., Bonnevie, T., Moser, M. B. & Moser, E. I. Hippocampus-independent phase precession in entorhinal grid cells. Nature 453, 1248–1252 (2008).

22. Qasim, S. E., Fried, I. & Jacobs, J. Phase precession in the human hippocampus and entorhinal cortex. Cell 184, 3242–3255.e10 (2021).

23. Moguilner, S. G. et al. Sleep functional connectivity, hyperexcitability, and cognition in Alzheimer’s disease. Alzheimer’s and Dementia 20, 4234–4249 (2024).

24. Oostenveld, R., Fries, P., Maris, E. & Schoffelen, J. M. FieldTrip: Open source software for advanced analysis of MEG, EEG, and invasive electrophysiological data. Comput Intell Neurosci 2011, (2011).

25. Berens, P. CircStat: A MATLAB Toolbox for Circular Statistics. J Stat Softw 31, 1–21 (2009).

