## Supplementary Information for "Phase precession of spindle-slow oscillation coupling across the human brain"

**eTable 1. Characteristics of Participants**

| | EMU | AMB | p-value |
|---|---|---|---|
| **N** | 58 | 43 | |
| **Age** | | | |
| **mean** | 45 | 73 | <0.001 |
| **SD** | 16.2 | 9.1 | |
| **range** | 20-80 | 51-85 | |
| **Sex (% Male)** | 26 | 44 | 0.08 |

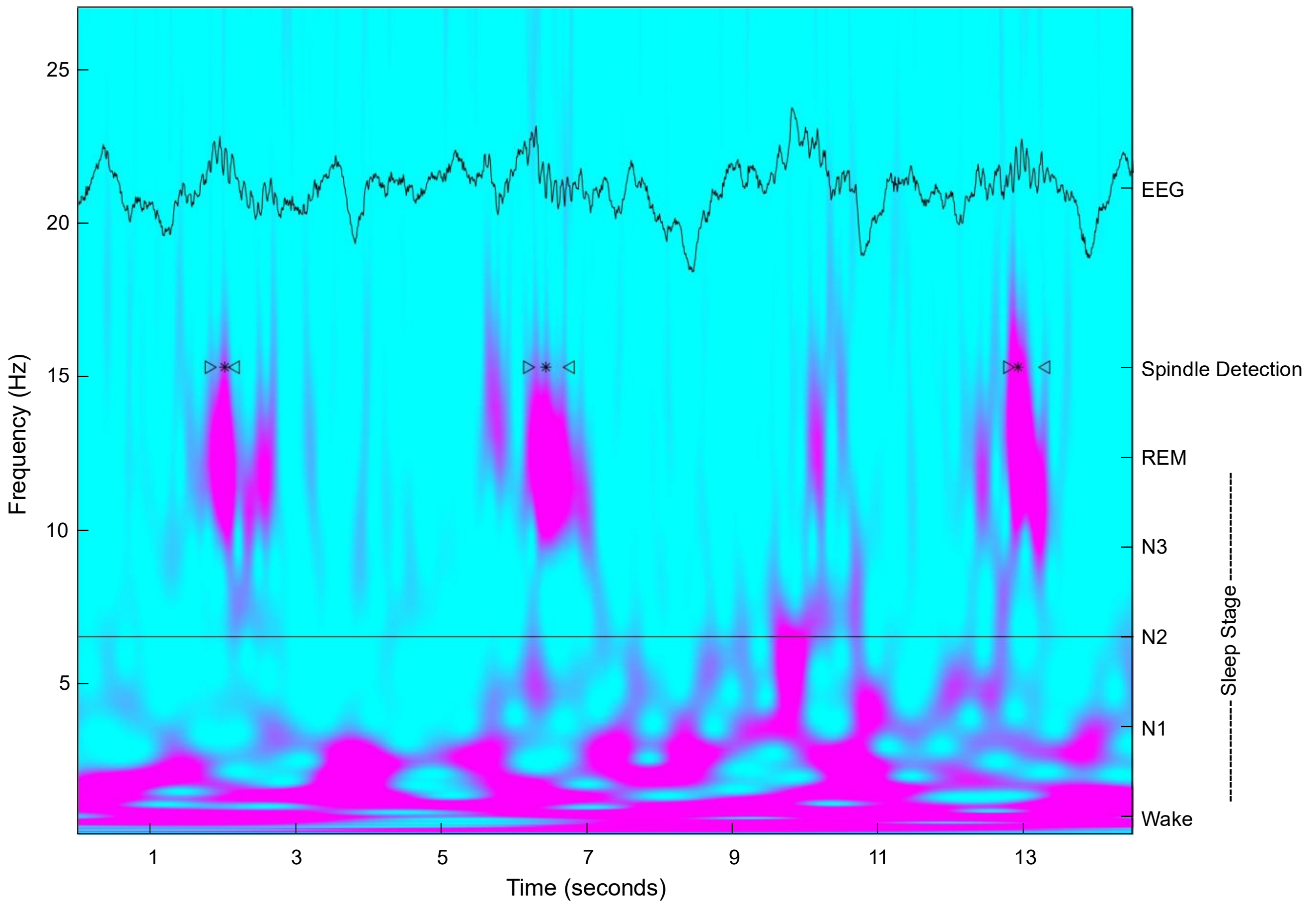


**eFigure 1. Spectral analysis of spindle and slow oscillations.** A time-frequency spectrogram is shown for 15 seconds of EEG in N2 sleep. The corresponding raw EEG tracing is superimposed (top). Automatic spindle detections are indicated by the >*< symbols. In this epoch, 3 automatically-detected spindles are seen, each occurring near the peak of the SO signal.

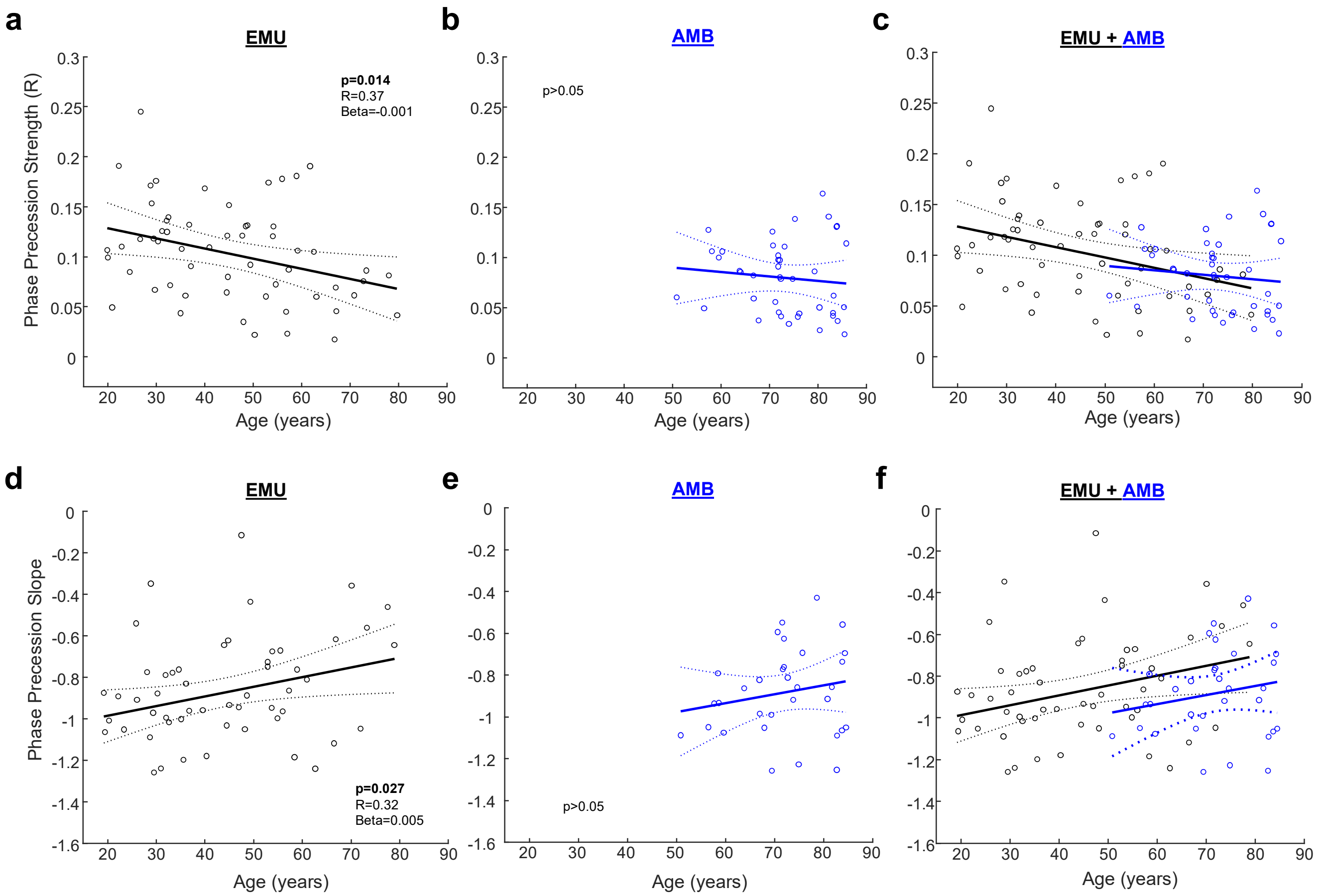


**eFigure 2. Sub-group analysis of the association of phase precession properties with age**. Data are shown for EMU and AMB participants separately. **a-c,** Scatter plot of model-adjusted phase precession strength versus age. There was a significant negative association in EMU participants. AMB participants, representing a narrower age range (from 50-85 years), did not demonstrate a significant association. Combined data are shown in **c** and in Fig. 4a. **d-f**, Scatter plot of model-adjusted phase precession slope versus age. There was a significant positive association in EMU, but not AMB participants. Combined data are shown in **f** and Fig 4c.
